# The native mussel *Mytilus chilensis* genome reveals adaptative molecular signatures facing the marine environment

**DOI:** 10.1101/2022.09.06.506863

**Authors:** Cristian Gallardo-Escárate, Valentina Valenzuela-Muñoz, Gustavo Nuñez-Acuña, Diego Valenzuela-Miranda, Fabian Tapia, Marco Yévenes, Gonzalo Gajardo, Jorge E. Toro, Pablo A. Oyarzún, Gloria Arriagada, Beatriz Novoa, Antonio Figueras, Steven Roberts, Marco Gerdol

## Abstract

The blue mussel *Mytilus chilensis* is a key socioeconomic species inhabiting the southern coast of Chile. This endemic marine mussel supports a booming aquaculture industry, which entirely relies on artificially collected seeds from natural beds that are translocated to a diverse physical-chemical ocean conditions’ for farming. Furthermore, mussel production is threatened by a broad range of microorganisms, pollution, and environmental stressors that eventually impact its survival and growth. Herein, understanding the genomic basis of the local adaption is pivotal to developing sustainable shellfish aquaculture. We present a high-quality reference genome of *M. chilensis*, which is the first chromosome-level genome for a Mytilidae member in South America. The assembled genome size was 1.93 Gb, with a contig N50 of 134 Mb. Through Hi-C proximity ligation, 11,868 contigs were clustered, ordered, and assembled into 14 chromosomes in congruence with the karyological evidence. The *M. chilensis* genome comprises 34,530 genes and 4,795 non-coding RNAs. A total of 57% of the genome contains repetitive sequences with predominancy of LTR-retrotransposons and unknown elements. Comparative genome analysis was conducted among *M. chilensis* and *M. coruscus*, revealing genic rearrangements distributed into the whole genome. Notably, Steamer-like elements associated with horizontal transmissible cancer were explored in reference genomes, suggesting putative phylogenetic relationships at the chromosome level in Bivalvia. Genome expression analysis was also conducted, showing putative genomic differences between two ecologically different mussel populations. Collectively, the evidence suggests that local genome adaptation can be analyzed to develop sustainable mussel production. The genome of *M. chilensis* provides pivotal molecular knowledge for the *Mytilus* complex evolution and will help to understand how climate change can impact mussel biology.

## 1. INTRODUCTION

The blue mussel *Mytilus chilensis* (Hupé, 1854) is an ecological and socioeconomic key species in Chile, that leads the national shellfish aquaculture. This endemic mussel species constitutes one of the main industries in mussel production worldwide (Gonzalez-Poblete, Hurtado, Rojo, & Norambuena, 2018; Uriarte, 2008). However, the success of mussel aquaculture production in Chile is threatened by a wide range of microorganisms (Detree, Nunez-Acuna, Roberts, & Gallardo-Escarate, 2016; Enriquez, Frosner, Hochsteinmintzel, Riedemann, & Reinhardt, 1992; Gray, Lucas, Seed, & Richardson, 1999), marine pollution (Blanc, Molinet, Subiabre, & Diaz, 2018) and the climate variability that can impact the larval settlement and growth of mussel populations (Harvell, 2002; Hüning, 2013; Vihtakari et al., 2013). To cope with those stressors, mussels and marine invertebrates produce two-component responses, a specific response to the stressor and a more general response involving immune and endocrine pathways (Cooper 1996). Impacts of multi-stressors have been predicted to have additive, synergetic or antagonistic effects on marine organisms’ physiology (Crain, Kroeker, & Halpern, 2008a). These different effects are directly linked to the amount of time between the occurrence of stressors, their intensity, and the organism’s ability to return to homeostasis before a new stressor occurs (Gunderson, Armstrong, & Stillman, 2016). Despite these predictions, meta-data analyses show that most of the studied multi-stressors had synergetic effects on organisms’ physiology (Crain, Kroeker, & Halpern, 2008b). Notably, isolated effects on mussel’s immune system of environmental stressors or pathogens infection have been extensively studied (Bibby, Widdicombe, Parry, Spicer, & Pipe, 2008; Lockwood, Sanders, & Somero, 2010; Malagoli, Casarini, Sacchi, & Ottaviani, 2007; Mitta et al., 2000; Pereiro, Moreira, Novoa, & Figueras, 2021; Rey-Campos, Novoa, Pallavicini, Gerdol, & Figueras, 2021; Romero, Novoa, & Figueras, 2022; Sendra et al., 2020). However, mussel’s immune response to the combination of stressors remains unexplored. The interplaying between the immune system and multi-environmental stressors such as ocean acidification, hypoxia, marine heatwaves, HABs, and pathogen infections requires physiological, cellular, and molecular tools that uncover the complexity of mussel biology. Here, high-quality genome assembly at the chromosome level is pivotal to driving the scientific community in this endeavor and mussels represent an outstanding model species. For instance, mussel species display morphologically conserved karyotypes, and there is no evidence of whole-genome duplication events (Perez-Garcia, Moran, & Pasantes, 2014). Compared with other bivalves, the reported mussel genomes share relatively large genome sizes characterized by high heterozygosity and expanded mobile elements (Du et al., 2017; McCartney et al., 2022; Uliano-Silva et al., 2017; S. Wang et al., 2017; Zhang et al., 2012). Unfortunately, these genome features challenge the assembly efforts to avoid genome fragmentation. Up to now, chromosome-level genome assembly for Mytilidae has only been reported for the congeneric species *M. coruscus* (Yang et al., 2021) and for the zebra mussel *Dreissena polymorpha* (McCartney et al., 2022); meanwhile, other members of the Family have been reported as highly contiguous reference assemblies at contig level (Murgarella et al., 2016; Renaut et al., 2018; Xu et al., 2017). Interestingly, the presence-absence variation (PAV) phenomenon has recently been reported for *M. galloprovincialis*, where a pan-genome composed of 20,000 protein-coding genes was observed in conjunction with dispensable genes that are entirely missing in some mussels (Gerdol et al., 2020).

The endemic Chilean blue mussel *M. chilensis*, a close relative of the *M. edulis* species complex of the northern hemisphere (Gaitan-Espitia, Quintero-Galvis, Mesas, & D’Elia, 2016; Larrain, Zbawicka, Araneda, Gardner, & Wenne, 2018), represents an iconic species to explore key questions in ecology (Curelovich, Lovrich, Cueto, & Calcagno, 2018), ecophysiology (Duarte et al., 2018) and adaptative genomics (Yevenes, Nunez-Acuna, Gallardo-Escarate, & Gajardo, 2021, 2022). It is a keystone taxon in the ecosystem regulating phytoplankton and nutrient flow and contributes to remineralizing organic deposits in the sediment (Hargrave, Doucette, Cranford, Law, & Milligan, 2008; Vinagre, Mendonca, Narciso, & Madeira, 2015). It inhabits rocky substrates in the intertidal and subtidal zones along the southern Pacific Ocean from latitude 38S to 53S (Flores et al., 2015). As a gonochoric species with an annual gametogenic cycle, sexual maturity occurs in spring-summer, where planktonic larvae can drift between 20 and 45 days before settlement (Toro, Ojeda, & Vergara, 2004). Dispersal potential has been estimated to be up to 30 km, allowing different degrees of gene flow among mussel populations (Astorga, Vargas, Valenzuela, Molinet, & Marin, 2018).

In this study, PacBio sequencing, and Hi-C scaffolding technology were jointly used to assemble the first chromosome-level reference genome of *M. chilensis*. Moreover, we conducted a comparative genomics study among reported genome mussel’s species and analyzed the molecular signatures in two mussel populations facing distinct physical-chemical ocean conditions. Genomic features revealed putative chromosome rearrangements among mussel species, suggesting phylogenetic relationships for Steamer-like elements in Mytilidae. Candidate genes and single nucleotide polymorphisms were also associated with local adaptation of *M. chilensis*, revealing specific-transcriptome profiles associated with metabolism and immune-related genes. The knowledge gained in this study will provide pivotal information to explore how the marine environment drives phenotypic plasticity, that in turn, reveals genome adaptation signatures in mussel populations.

## 2. MATERIAL AND METHODS

### 2.1. Sample collection, NGS libraries, and sequencing

Adult specimens of *M. chilensis* were collected from a natural bed located in Puerto Mont (41°48’S-73°5’W), Chile (Figure 1A). Five mussels were selected for whole-genome sequencing using 1 mL of hemolymph collected from each specimen to reduce the heterozygosity or the number of individuals per pool. The samples were centrifuged at 1200 RPM to isolate the hemocyte cells and preserved by liquid nitrogen. High-quality DNA was isolated using the Qiagen DNA purification kit (QIAGEN, Germantown, MD, USA) following the manufacturer′s instructions and quantified by TapeStation 2200 instrument (Agilent, USA). DNA samples >9.5 in DIN numbers were selected for library preparation. Furthermore, 50 individuals were sampled from Cochamó (41°28’S-72ª18’W) and Yaldad (43°07’S-73°44’W), Southern Chile, to isolate RNA and explore molecular signatures associated with the local genome adaptation. Herein, these mussel populations inhabit contrasting ocean variability characterized by an estuary with continuous input of freshwater and vertical stratification and a bay exposed to open sea influence, respectively. Over the past two decades, the temporal and spatial variability of Sea Surface Temperature (SST) around Puerto Montt, Chiloé island and at sites Yaldad and Cochamó were analyzed using satellite images. Data on sea surface temperature in the region of interest were obtained from MUR-SST (Multi-scale Ultra-High-Resolution SST) distributed by NOAA through its ERDDAP platform. The MUR-SST images have a spatial resolution of 1 km and a temporal resolution of 1 day. *In situ* measurements for temperature (°C) and salinity (PSU) seawater were obtained for both locations between 2018 and 2019. The raw environmental data were collected from the CHRONOS database, managed by Instituto de Fomento Pesquero, IFOP (Chile).

**Figure 1.**
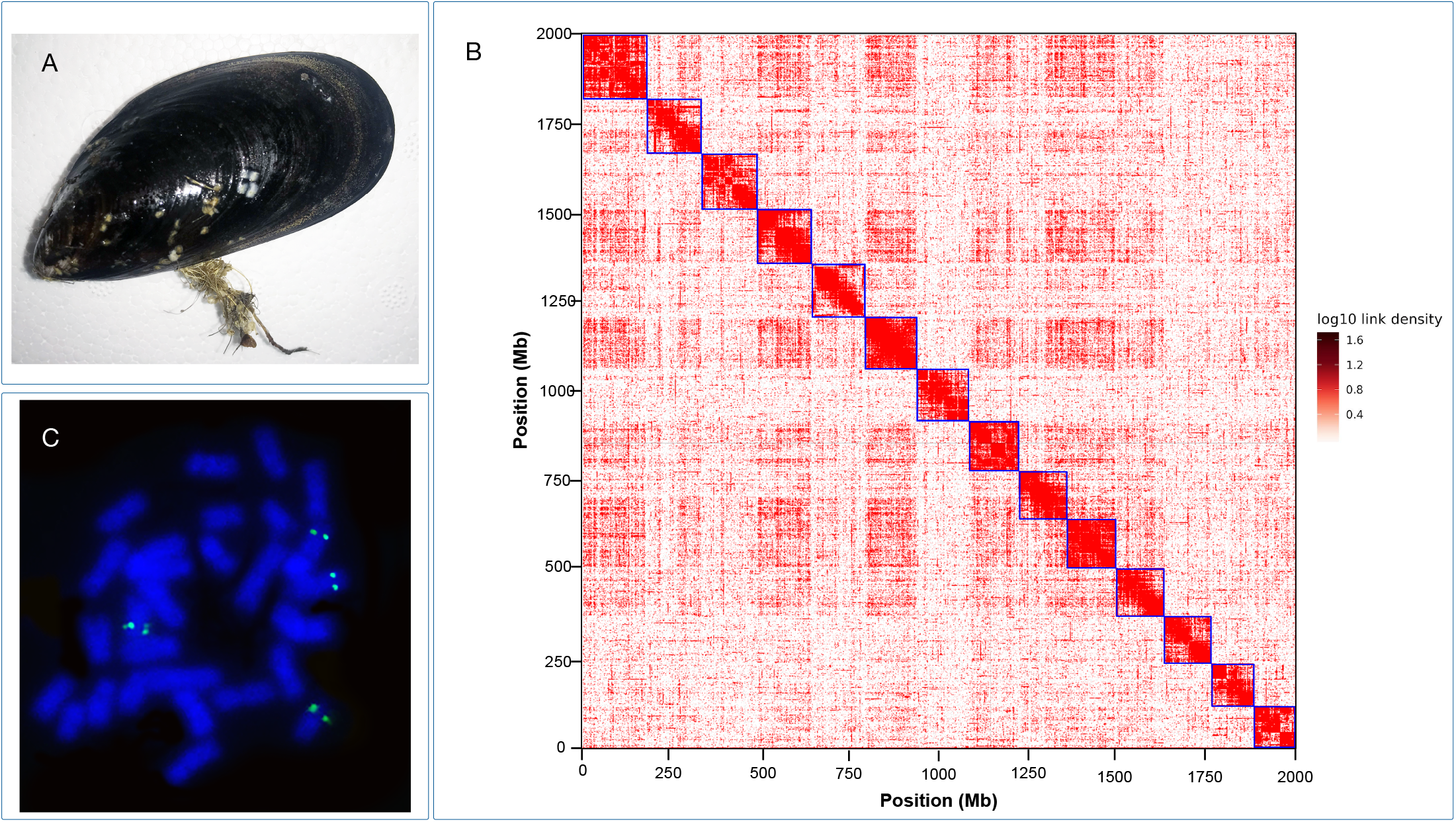
(A) Photograph of the native blue mussel *Mytilus chilensis*. (B) Metaphasic chromosomes from mussel larvae samples and mapping of 28S-rDNA by fluorescence *in situ* hybridization. (C) Heatmap of chromosome interaction intensity in the blue mussel Hi-C assembly. The x-and y-axis represents the length of the chromosomes. The color bar represents the Hi-C contact density.

Samples were prepared according to the SMRTbell guide for sequencing on the PacBio Sequel II System. The genomic DNA isolated from 5 individuals collected from Puerto Montt was sequenced using SMRT Sequencing according to the manufacturer’s protocols. SMRT sequencing yielded 882.1 Gigabases and 63 million long reads from 2 HiFi SMRT cells. The subreads N50 and average read lengths were 14,665 and 14,535 bp, respectively. The total HiFi reads yielded 3.7 million with an average quality of Q36 and Q35, respectively. Hi-C libraries were constructed from hemocyte cells using Phase Genomics’ Animal Hi-C kit and sequenced on Illumina’s Hiseq4000 platform to yield 253 million reads using the same DNA isolated for PacBio sequencing. Short-read sequencing libraries were prepared using an insert size of 150 bp obtained from 1 μg of genomic DNA after fragmentation, end-paired, and ligation to adaptors. The ligated fragments were fractionated on agarose gels and purified by PCR amplification to produce sequencing libraries. The method applied was like that previously published by Lieberman-Aiden et al. (2009). The PacBio and Hi-C Illumina DNA raw data were deposited in the Sequence Read Archive (SRA) repository, accession number SRR20593343 and SRR20966976, respectively.

Moreover, RNA libraries were constructed from hemocytes, digestive gland, gill, and mantle tissues for transcriptome sequencing to obtain whole-transcriptome profiling from the same mussels used for genome DNA sequencing. Additionally, twelve available Sequence Read Archive (SRA) transcriptomic data (GenBank accession number SRP261955), representing gills and mantle tissues collected from individuals of Cochamó and Yaldad mussel populations (Yévenes et al., 2021, 2022), were incorporated to analyze population-specific transcriptome profiles. These twelve transcriptomic SRA data represent to total RNA extracted by the Trizol reagent method (Invitrogen, USA) from 15 individuals. Three biological replicates (5 individual total RNA extractions each one) from each mussel population were analyzed. The quality and integrity of extracted RNAs were measured in a TapeStation 2200 instrument (Agilent, USA), using the R6K Reagent Kit based on the manufacturer’s instructions. RNA samples >9 in RIN numbers were selected for high-quality libraries preparation using TrueSeq Stranded mRNA LT Sample Prep Kit and sequenced in HiSeq 4000 (Illumina, USA).

### 2.2. *De novo* genome assembly and Hi-C scaffolding of *M. chilensis*

Two HiFi single-molecular real-time cells in the PacBio Sequel platform yielded 53.8 Gb of high-quality DNA genome information. These long reads were assembled with the Hifiasm package using default parameters (Cheng, Concepcion, Feng, Zhang, & Li, 2021). For Hi-C scaffolding, reads were aligned to the primary draft assembly, also following the manufacturer’s instructions (Phase-Genomics, 2019). Briefly, reads were aligned using BWA-MEM (H. Li & Durbin, 2010) with the –5SP and –t 8 options specified, and all other options default. The package SAMBLASTER (Faust & Hall, 2014) was used to flag duplicates and then excluded for further analysis. Sequence alignments were filtered with SAMtools (Danecek et al., 2021; H. Li et al., 2009) using the –F 2304 filtering flag to remove non-primary and secondary alignments. This step was conducted to remove alignment errors, low-quality alignments, and other alignment noise due to repetitiveness, heterozygosity, and other ambiguous assembled sequences. Finally, Phase Genomics’ Proximo Hi-C genome-scaffolding platform was used to create chromosome-scale scaffolds from the corrected assembly, according to Bickhart et al. (2017).

### 2.3. Karyotype of *M. chilensis*

Metaphase plates of 24 hours post-fertilization larvae were used to obtain chromosomes from *M. chilensis*, according to Gallardo-Escárate et al. (2004). Briefly, antimitotic treatment with Colchicine 0.05% solution was applied for 4 hours. Then the larvae were rinsed in clean seawater and immersed in a hypotonic solution (seawater: distilled water, 1:1) for 30 min. Finally, the larvae were fixed in modified Carnoy solution (methanol: acetic acid, 3:1). Chromosome spreads were obtained by dissociating larva tissue in acetic acid (50%), pipetting suspension drops onto slides preheated at 43°C and air-dried according to Amar et al. (2008). FISH experiment was performed to validate the physical localization of candidate genes. Here, 28S rDNA was labeled following methods previously published (Perez-Garcia et al., 2014). Briefly, metaphase preparations were denatured at 69°C for 2 min and hybridized overnight at 37°C. Signal detection was performed using fluorescein avidin and biotinylated anti-avidin for the biotinylated probes and mouse anti-digoxigenin, goat anti-mouse rhodamine, and rabbit anti-goat rhodamine for the digoxigenin-labeled probes. Fluorescent staining was carried out with 4,6-diamidino-2-phenylindole (DAPI) and mounted with Vectacshield antifading solution. Chromosome spreads were observed using an epifluorescent microscope Nikon Eclipse 80i equipped with a digital camera DS-5Mc.

### 2.4. Genome annotation of *M. chilensis*

Our repeat annotation pipeline applied a combined homology alignment strategy, and *de novo* search to identify the whole genome repeats. Tandem Repeat was extracted using TRF (http://tandem.bu.edu/trf/trf.html) by *ab initio* prediction. The homolog prediction commonly used Repbase (www.girinst.org/repbase) database employing RepeatMasker (www.repeatmasker.org/) software and its in-house scripts (RepeatProteinMask) with default parameters to extract repeat regions. *Ab initio* prediction was used to build a de novo repetitive elements database by LTR_FINDER (http://tlife.fudan.edu.cn/ltr_finder/), RepeatScout (www.repeatmasker.org/), RepeatModeler (www.repeatmasker.org/RepeatModeler.html) with default parameters. Then, all repeat sequences with lengths >100bp and gap ‘N’ less than 5% constituted the raw transposable element (TE) library. A custom library (a combination of Repbase and a custom *de novo* TE library processed by UCLUST to yield a non-redundant library) was supplied to RepeatMasker for DNA-level repeat identification.

The structural annotation approach was applied to incorporate *de novo*, homolog prediction, and RNA-Seq-assisted predictions to annotate gene models. For gene prediction based on *de novo*, Augustus (v3.2.3), Geneid (v1.4), Genescan (v1.0), GlimmerHMM (v3.04) and SNAP (2013-11-29) were used in our automated gene prediction pipeline. For homolog prediction, sequences of homologous proteins were downloaded from Ensembl/NCBI/others. Protein sequences were aligned to the genome using TblastN (v2.2.26; E-value ≤ 1e−5), and then the matching proteins were aligned to the homologous genome sequences for accurate spliced alignments with GeneWise (v2.4.1) software to predict gene structure contained in each protein region. Finally, for RNA-seq data, transcriptome reads assemblies were generated with Trinity (v2.1.1) for the genome annotation. For the genome annotation optimization, the RNA-Seq reads from different tissues were aligned to genome fasta using Hisat (v2.0.4)/TopHat (v2.0.11) with default parameters to identify exons region and splice positions. The alignment results were inputted for Stringtie (v1.3.3)/Cufflinks (v2.2.1) with default parameters for genome-based transcript assembly. The non-redundant reference gene set was generated by merging genes predicted by three methods with EvidenceModeler (EVM v1.1.1) using PASA (Program to Assemble Spliced Alignment) terminal exon support and including masked transposable elements as input into gene prediction. Individual families of interest were selected for further manual curation.

Gene functions were assigned according to the best match by aligning the protein sequences to the Swiss-Prot using Blastp (with a threshold of E-value ≤ 1e−5). The motifs and domains were annotated using InterProScan70 (v5.31) by searching against publicly available databases, including ProDom, PRINTS, Pfam, SMRT, PANTHER, and PROSITE. The Gene Ontology (GO) IDs for each gene were assigned according to the corresponding InterPro entry. We predicted the protein function by transferring annotations from the closest BLAST hit (E-value<1e−5) in the Swissprot database and DIAMOND (v0.8.22) / BLAST hit (E-value<1e−5) in the NR database. We also mapped the gene set to a KEGG pathway and identified the best match for each gene.

Non-coding RNA annotations such as tRNAs were predicted using the program tRNAscan-SE (http://lowelab.ucsc.edu/tRNAscan-SE/. Since rRNAs are highly conserved, we choose relative species’ rRNA sequences as references and predict rRNA sequences using Blast. Other ncRNAs, including miRNAs and snRNAs, were identified by searching against the Rfam database with default parameters using the infernal software (http://infernal.janelia.org/). Additionally, lncRNAs were identified using the pipelines previously proposed (Gallardo-Escarate, Figueras, & Novoa, 2019; Pereiro et al., 2021; Tarifeno-Saldivia, Valenzuela-Miranda, & Gallardo-Escarate, 2017).

### 2.5. Comparative genomics between *M. chilensis* and *M. coruscus*

Syntenic relationships were carried out among mussel species for which chromosome-level reference genomes are publicly available. Here, the analysis was performed between the two congeneric species *M. chilensis* (this study) and *M. coruscus* (Yang et al., 2021), where gene annotations were explored by MCScanX (Y. P. Wang et al., 2012) implemented in the TBtools package (Chen et al., 2020). This approach detects groups of orthologous genes and compares their arrangement to identify colinear segments in the compared genomes. MCScanX was used to discover microsynteny relationships, focusing on the local arrangement of genes near the syntenic blocks. The microsynteny arrangement of genes identified by MCScanX was evaluated through GO analysis to identify the primary molecular function and biological processes enrichened for each genomic region where macromutations or chromosome rearrangements were detected.

Disseminated neoplasia is a disease horizontally transmitted by clonal cancer cells, which causes leukemia in mollusk bivalves (Metzger, Reinisch, Sherry, & Goff, 2015; Metzger et al., 2016). The neoplastic cells gradually replace normal hemocytes leading to increased mortality, and it has been detected in 15 species of marine bivalve mollusks worldwide (Elston, Moore, & Brooks, 1992). Notably, disseminated neoplasia has been observed among mussel species with varying epizootic prevalences. For instance, *M. trossulus* has shown high prevalences in some areas, whereas in *Mytilus edulis*, the prevalences are generally lower. Furthermore, *M. galloprovincialis* has been suggested as a resistant species to the disease in Spain and Italy’s mussel populations (Ciocan & Sunila, 2005). This observation extends the relevance to exploring mussel species’ genetic features associated with disseminated neoplasia. Herein, the molecular characterization of steamer-like elements in *M. chilensis* was conducted by cloning and walking primer method according to Arriagada et al. (2014). The putative *M. chilensis* Steamer-like was scanned through twelve reference genomes assembled at chromosome level for Bivalvia: *Mytilus coruscus* (GCA_017311375.1), *Mytilus edulis* (GCA_019925275.1), *Dreissena polymorpha* (GCA_020536995.1), *Mercenaria mercenaria* (GCA_014805675.2), *Solen Grandis* (GCA_021229015.1), *Ruditapes philippinarum* (GCA_009026015.1), *Pecten maximus* (GCA_902652985.1), *Pinctada imbricata* (GCA_002216045.1), *Crassostrea gigas* (GCA_902806645.1), *Crassostrea ariakensis* (GCA_020458035.1), and *Crassostrea virginica* (GCA_002022765.4). The putative Long Terminal Repeat (LTR) sequences were identified using BLAST search, where open reading frames (ORFs) between flanking LTRs were detected. The identified Steamer-like elements were aligned using ClustalW and annotated based on NCBI Conserved Domain search. Amino acid sequences for the full-length Gag-Pol polyprotein region were aligned among the studied bivalve species. The Steamer element was reported for *Mya arenaria* (Accession AIE48224.1) and *M. chilensis* (this study). DNA sequence genealogy analysis was conducted to investigate horizontal transmission events among bivalve species. The maximum likelihood (ML) method was conducted on the SLEs loci localized in all the publicly available bivalve genomes assembled at the chromosome level.

### 2.6. Whole-genome transcript expression analysis in two *M. chilensis* populations

The transcriptome of mussels collected from Yaldad and Cochamó populations were analyzed using an hierarchical clustering approach to detect transcriptional similarities among tissues/populations. The transcripts that were differentially expressed in comparison to normalized expression values were visualized in a clustering heat map and selected according to the identified cluster. For an optimal comparison of the results, k-means clustering was performed to identify candidate genes involved in specific gene expression patterns. The distance metric was calculated with the Manhattan method, where the mean expression level in 5-6 rounds of k-means clustering was subtracted.

Moreover, raw data from mussel tissues collected from Yaldad and Cochamó populations were trimmed and mapped to the *M. chilensis* genome using CLC Genomics Workbench v22 software (Qiagen Bioinformatics, USA). Threshold values for transcripts were calculated from the coverage analysis using the Graph Threshold Areas tool in CLC Genomics Workbench v22 software. Here, an index denoted as Chromosome Genome Expression (CGE) was applied to explore the whole-genome transcript expression profiling, according to Valenzuela et al. (2022). The CGE calculates the mean coverage of transcripts mapped into a specific chromosome region, comparing mussel populations and tissues. Specifically, the CGE index represents the percentage of the transcriptional variation between two or more RNA-seq data for the same locus. The transcript coverage values for each dataset were calculated using a threshold of 20,000 to 150,000 reads, where a window size of 10 positions was set to calculate and identify chromosome regions differentially transcribed. This approach was used to visualize chromosome regions actively transcribed, identifying genes differentially expressed and observing tissue-specific patterns in the evaluated mussel populations. Finally, the threshold values for each dataset and CGE index were visualized in Circos plots (M. I. Krzywinski et al., 2009).

RNA-seq data analyses were carried out using the raw sequencing reads and mapped on the assembled genome by CLC Genomics Workbench v22 software (Qiagen Bioinformatics, USA) for each tissue/population separately. In parallel, *de novo* assembling was performed to evaluate PAVs and dispensable genes affecting the *in-silico* transcription analysis. The assembly was performed with overlap criteria of 70% and a similarity of 0.9 to exclude paralogous sequence variants. The settings used were set as mismatch cost = 2, deletion cost = 3, insert cost = 3, minimum contig length = 200 base pairs, and trimming quality score = 0.05 using CLC Genomics Workbench v22. After assembly, the contigs generated for each data set were mapped on the genes annotated in the reference genome to evaluate de genome coverage and detect PAV features. The analysis did not show bias putatively associated with PAVs between the analyzed mussel populations. Then, mRNA sequences annotated for the *M. chilensis* genome were used to evaluate the transcription level between mussel populations, where differentially expression analysis was set with a minimum length fraction = 0.6 and a minimum similarity fraction (long reads) = 0.5. The obtained genes from each tissue/population were blasted to CGE regions to enrich the number of transcripts evaluated by RNA-Seq analysis. In addition, sequences were extracted near the threshold areas in a window of 10 kb for each transcriptome. The expression value was set as transcripts per million model (TPM). The distance metric was calculated with the Manhattan method, with the mean expression level in 5-6 rounds of k-means clustering subtracted. Finally, Generalized Linear Model (GLM) available in the CLC software was used for statistical analyses and to compare gene expression levels in terms of the log_2_ fold change (p = 0.005; FDR corrected). Moreover, the innate immunity in marine invertebrates may play an important role in speciation and environmental adaptation (Ellis et al., 2011; Rolff & Siva-Jothy, 2003). Herein, we investigate the immune-related genes associated with the TOLL-like receptor (TLR) and Apoptosis signaling pathways due that the functional annotation revealed thar they were mainly enriched between the mussel populations analyzed. In addition, bioinformatic analyses were carried out using the CLC Genomics Workbench software to mine single nucleotide variants (SNV) from the transcriptomes sequenced for Yaldad and Cochamó. Candidate SNVs were called with the following settings: window length = 11, maximum gap and mismatch count = 2, minimum average quality of surrounding bases = 15, minimum quality of central base = 20, maximum coverage = 100, minimum coverage = 8, minimum variant frequency (%) = 35.0, and maximum expected variations (ploidy) = 2. Furthermore, the genotypes of DEGs were also identified for detecting putative genetic variations between mussel populations. Here, singleton, dispersed, tandem, proximal, and whole genome duplication (WGD) gene events were evaluated using MCScanX. The amino acid changes and the zygosity proportions were also estimated in DEGs between Yaldad and Cochamó populations.

### 2.7. GO enrichment analysis

Differentially expressed mRNA were annotated through BlastX analysis using a custom protein database constructed from GeneBank, KEGG, GO, and UniProtKB/Swiss-Prot. The cutoff E-value was set at 1E-10. Transcripts were subjected to gene ontology (GO) analysis using the Blast2GO plugins included in the CLC Genomics Workbench v22 software (Qiagen Bioinformatics, USA). The results were plotted using the Profiler R package (Yu, Wang, Han, & He, 2012). GO enrichment analysis was conducted to identify the most represented biological processes among protein-coding genes proximally located to the CGE regions. The enrichment of biological processes was identified using Fisher’s exact test tool of Blast2GO among the different tissues and mussel populations.

## 3. RESULTS AND DISCUSSION

### 3.1. Chromosome genome assembly of *M. chilensis* using proximity ligation

With two HiFi single-molecular real-time cells in the PacBio Sequel platform, we generated 53.8 Gb of high-quality DNA genome information. This data comprised 63 million reads with a total length of 882 Giga base pairs (Table1). These long reads were assembled with the Hifiasm package using default parameters (Cheng et al., 2021), yielding a primary assembly of 13,762 contigs equivalent to 2.19 Gb, with an N50 of 206 Mb. The size genome assembly made by Hifiasm was comparable with the previous genome size described for closely related species; 1.28 Gb for *M. galloprovincialis* (Gerdol et al., 2020), 1.57 Gb for *M. coruscus* (Yang et al., 2021) and 1.79 Gb for *Dreissena polymorpha* (McCartney et al., 2022).

*In vivo* Hi-C is a technique that maps physical DNA-DNA proximity across the entire genome (Ghurye et al., 2019; Pal, Forcato, & Ferrari, 2019). The method was introduced as a genome-wide version of its predecessor, 3C (Chromosome Conformation Capture). It has been a powerful tool in chromosome-scale genome assembly of many animals in recent years (Lando, Stevens, Basu, & Laue, 2018; Lin et al., 2018). In this study, Hi-C experiments and data analysis on hemocyte cells were used for the chromosome assembly of the blue mussel *M. chilensis*. Here, two Hi-C libraries were prepared and sequenced by Phase Genomics (Seattle, WA, USA), resulting in ∼20x coverage and ∼253 million 150-bp paired-end reads (Table 1). The Hi-C analysis evidenced that 44.68% of high-quality reads showed intercontig signals or Cis-close position (<10kbp on the same contig), and an additional 4.09% of sequence reads revealed a Cis-far conformation (>10Kbp on the same contig) (Table 2). Hi-C reads were aligned using Bowtie (Langmead, Trapnell, Pop, & Salzberg, 2009) to order and orient the 13,762 contigs, and scaffolding was performed using Proximo (Phase Genomics, Seattle, WA, USA). We then applied Juicebox for visual inspection and manual correction (Robinson et al., 2018). We also manually removed 1,894 scaffolds that were microbe-sized and disconnected from the rest of the assembly. Herein, 11,868 contigs were used for the first chromosome-level high-quality *M. chilensis* assembly (Table 3). The N50 and total genome length were calculated in 134 Mbp and 1,938 Gbps, respectively. The *M. chilensis* genome provides a useful genomic resource for research in mussel biology and for developing novel sustainable strategies in mussel aquaculture. The Hi-C data generated 14 chromosomes assembled with HiFi consensus long DNA reads (Fig. 1B). The cytogenetic analysis performed for *M. chilensis* revealed a conservative karyotype for the *Mytilus* genus composed by 2n=14 (Perez-Garcia et al., 2014). Physical localization of 28S-rRNA revealed two loci mapped in different submetacentric/subtelocentric chromosome pairs (Fig. 1C), confirming the presence of major rDNA clusters subterminal to the long arms of two chromosome pairs reported in *M. edulis* and *M. galloprovincialis* (Martínez-Lage, González-Tizón, & Méndez, 1995). Concerning genome assembly, the largest scaffold was assembled from 998 contigs in a total size of 173.3 Mb. Meanwhile, the smallest scaffold was 117.3 Mb in length, consisting of 744 contigs (Table 3). Notably, the number of contigs in scaffolds were 11,868 (100% of all contigs in chromosome clusters, 86.24% of all contigs) and 1.93 Gbps of genome size (100% of all length in chromosome cluster, 88.43% of all sequence length). The completeness of genome assembly was assessed by the single-copy ortholog set (BUSCO, V5.3.2) (Manni, Berkeley, Seppey, Simao, & Zdobnov, 2021). The results showed the following BUSCO scores: i) Eukaryota Odb10; C:94.1% [S:72.9%, D:21.2%], F:3.1%, M:2.8%, n:255. ii) Metazoa Odb10; C:95.1% [S:75.5%, D:19.6%], F:2.5%, M:2.4%, n:954. iii) Mollusca Odb10, C:85% [S:70.1%, D:14.9%], F:3.6%, M:11.4%, n:5295.

**Table 1.** Statistics of whole-genome sequence assembly and transcriptome analysis of the blue mussel *Mytilus chilensis* using Illumina, PacBio, and Hi-C.

| Types | Method | No. of reads (Millions) | Library size | Length (Giga base pair) |
| --- | --- | --- | --- | --- |
| Genome | PacBio SMRT | 63M | 20kb | 882Gbp |
|  | Illumina (Hi-C) | 253M | 150bp | 37Gbp |
| Transcriptome | Illumina (Hemolymph) | 38.3M | 100bp | 3,838Gbp |
|  | Illumina (Mantle) | 37.1M | 100bp | 3,714Gbp |
|  | Illumina (Gills) | 38.6M | 100bp | 3,865Gbp |
|  | Illumina (Digestive gland) | 28.2M | 100bp | 2,823Gbp |

**Table 2.** Statistics of genome assembly using HiFi reads and proximity ligation analysis for *Mytilus chilensis*.

| Label | Statistics |
| --- | --- |
| ***PacBio assembly*** |  |
| Assembly size | 2,191,715,088 |
| Contig (CTG) N50 | 206,083 |
| CTGs | 13,762 |
| ***Hi-C mapping*** |  |
| Total read pairs (RPs) analyzed | 253,342,981 |
| High quality (HQ)* RPs | 14,71% |
| Clustering usable HQ reads per contig (CTGs >5KB)* | 1215.84 |
| RPs >10KB apart (CTGs >10KB) | 18.68% |
| Intercontig HQ RPs | 44.68% |
| Same strand HQ RPs | 21.50% |
| Split reads | 37.80% |

**Table 3.** *De novo* assembly of *M. chilensis* genome using proximity ligation (Hi-C).

| **Scaffold number** | **Number of contigs** | **Length (bp)** |
| --- | --- | --- |
| 1 | 988 | 173,300,526 |
| 2 | 852 | 140,500,440 |
| 3 | 957 | 154,573,458 |
| 4 | 871 | 155,184,769 |
| 5 | 902 | 143,621,794 |
| 6 | 845 | 146,028,403 |
| 7 | 946 | 139,985,977 |
| 8 | 821 | 134,004,722 |
| 9 | 794 | 133,130,516 |
| 10 | 821 | 132,801,123 |
| 11 | 802 | 131,378,307 |
| 12 | 790 | 122,972,572 |
| 13 | 735 | 113,313,251 |
| 14 | 744 | 117,342,092 |
| **Total** | **11,868** | **1,938,137,950** |
| N50 |  | 134,004,722 |

\*Number of contigs in scaffolds: 11,868 (100% of all contigs in chromosome clusters, 86.24% of all contigs)

### 3.2. Genome annotation of *M. chilensis*

The genome assembly was annotated using *de novo* and protein and transcript-guided methods (Fig. 2A). The first step of the annotation process was to identify the DNA repeats through the *M. chilensis* genome. Repetitive elements and non-coding genes in the blue mussel genome were annotated by homologous comparison and *ab initio* prediction. RepeatMasker (Bedell, Korf, & Gish, 2000) was used for homologous comparison by searching against the Repbase database (Bao, Kojima, & Kohany, 2015) and RepeatModeler (Storer, Hubley, Rosen, Wheeler, & Smit, 2021). According to these analyses, about 1.1 Gbps of repeat sequences were annotated, which accounted for 56.73% of the whole genome. Herein, DNA transposons, LINE, and LTR transposable elements were identified (Table 4). Useful genome information for population genetic studies is the identification of simple sequence repeats (SSRs) or microsatellites. The mining of SSRs revealed that the *M. chilensis* genome has 548,360 SSR sequences, where the 9% and 6% of the SSR loci were annotated for each mussel’s chromosome (Fig. 1S). The most frequent SSR motif was the tetranucleotide, followed by the dinucleotides, accounting for the 206,103 and 197,700 repeats, respectively. The entire SSR sequences accounted for 0.35% of the whole genome. The development of SSR markers offers a shortcut to assessing genetic diversity, which can potentially be applied in food authentication and genetic traceability for mussel species (Ferreira et al., 2020; Larrain, Diaz, Lamas, Uribe, & Araneda, 2014; Vidal, Penaloza, Urzua, & Toro, 2009).

**Figure 2.**
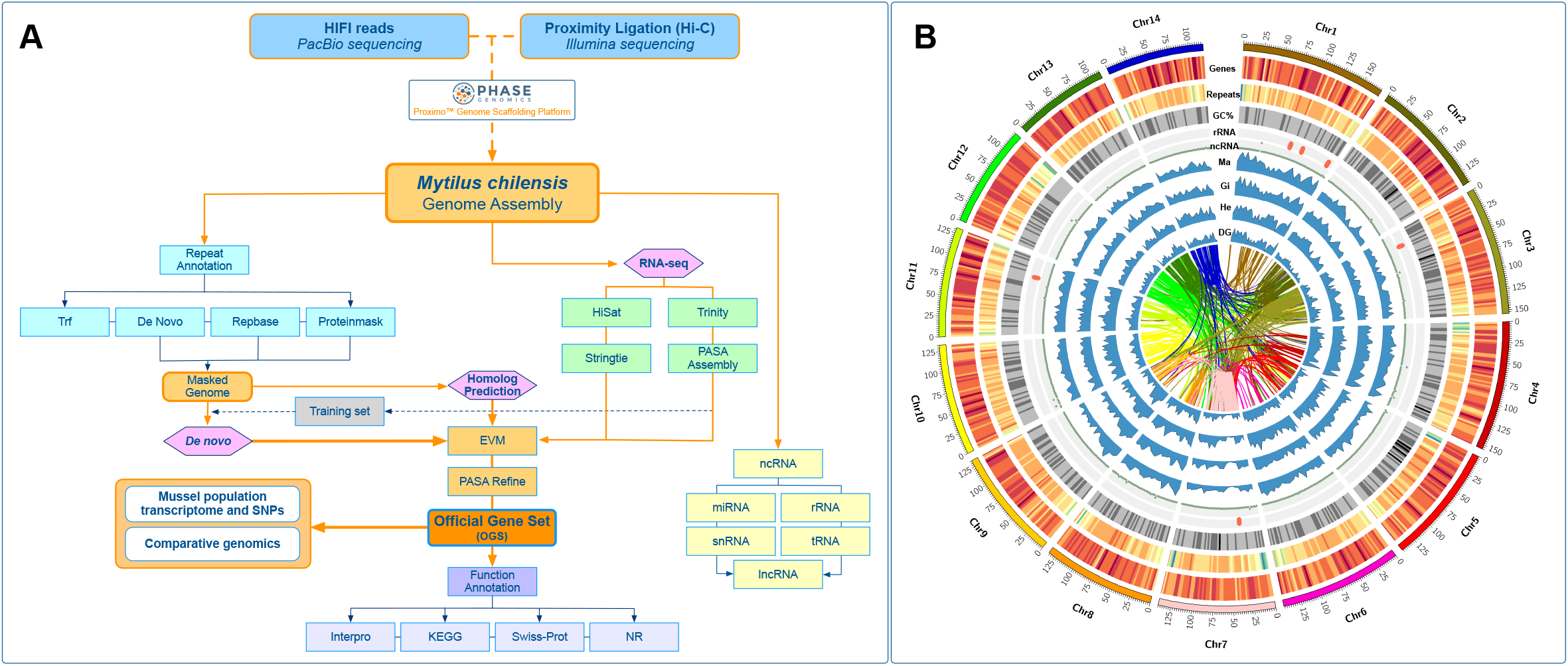
The native blue mussel *Mytilus chilensis* genome. (A) Workflow of *de novo* whole-genome sequencing project and annotation for *M. chilensis*. The rectangles indicate the steps of the primary data processing, and the arrows indicate output or input data. Pink diamonds indicate the combined strategy based on homolog prediction, de novo and RNA-seq assisted prediction. (B) The circos plot shows the genomic features for the 14 pseudo-chromosomes. From outer to the inner circle: Gene density, Repeat density, GC content, rRNAs localization and ncRNAs. The transcriptome expression for mantle (Ma), gills (Gi), Hemocytes (He), and digestive gland (DG) are shown as blue light profiles. Chromosome syntenies are represented in different colors according to each ideogram. The chromosome size is shown in the Mb scale.

**Table 4.**
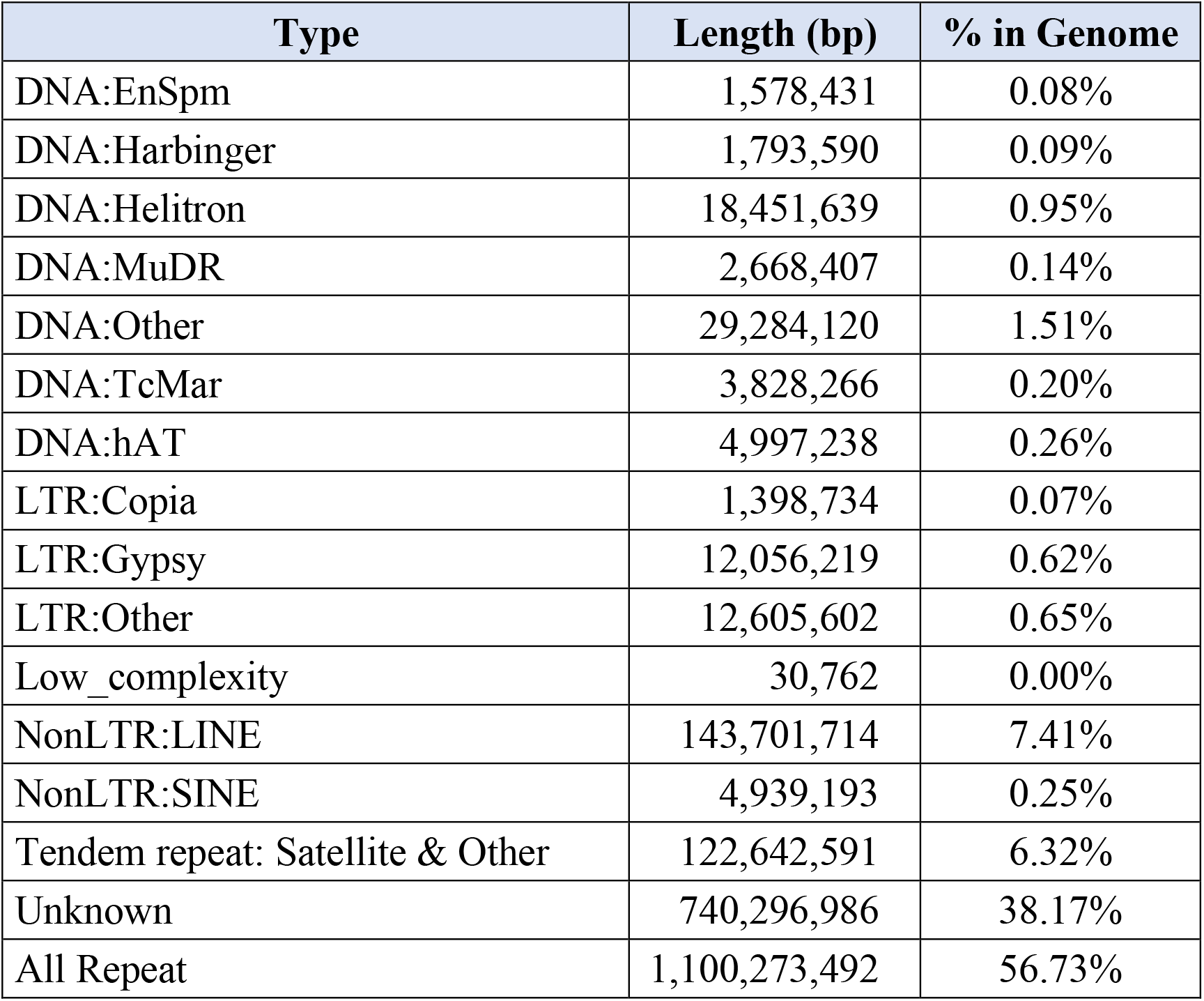
Statistics of the classification results of repeat sequence from *M. chilensis* genome

### 3.3. Protein-coding genes prediction and functional annotation in the *M. chilensis* genome

For the identification of protein-coding genes, *de novo*, homolog prediction, and RNA-seq evidence were used as the training set (Fig. 2A). For homologous predictions, the protein sequences from *Crassostrea gigas*, *Mytilus galloprovincialis*, *M. coruscus*, and *Dreissena polymorpha* genomes were extracted using the respectively published references and aligned against the blue mussel genome using TBLASTN (e-value<1e-5). The gene sequence structure of each candidate gene and previously mentioned tools were used to predict protein-coding genes. Finally, a non-redundant reference gene set was generated using EvidenceModeler (EVM) and PASA2 tools (Fig. 2A). Taken together, 34,530 protein-coding genes were identified with 6,531 bp of average transcript length, 1,377 bp of average CDS length, 4.92 of average of Exons per Gene, and 1,377 and 1,316 of the average length of exons and introns, respectively (Table 6). Additionally, 516 tRNAs were predicted using tRNAscan-SE, and 143 rRNA genes were annotated using RNAmmer. For non-coding RNAs with putative regulatory roles, 1,365 miRNAs and 43,011 long non-coding RNAs were identified and annotated within the *M. chilensis* genome (Table 7). For functional annotation, the predicted proteins within the blue mussel genome were searched by homology against seven databases: Swissprot, Nr, Nt, KEGG, eggnog, GO, and Pfam (Fig. 2A). Overall, 70.45%, 73.01%, 8.98%, 64.94%, 80.57%, 33.61% and 176.33 % of genes matched entries in these databases, respectively. A total of 34,530 genes (100%) were successfully annotated by gene function and conserved protein motifs (Table 8). The genomic features annotated for the native blue mussel *M. chilensis* were displayed using a circus plot (M. Krzywinski et al., 2009). Herein, this graphical representation showed the primary genomic features for the 14 chromosomes. Specifically, gene density, repeat density, GC content, rRNA localization, and ncRNAs were plotted. The transcriptome expression profiles for the mantle, gills, hemocytes, and digestive gland tissues were also displayed in connection with the syntenic blocks (Fig. 2B).

**Table 5.** Basic statistical results of gene structure prediction of relative species.

| Species | Number of genes | Average Transcript length (bp) | Average CDS length (bp) | Average Exons per gene | Average Exon length (bp) | Average Intron length (bp) |
| --- | --- | --- | --- | --- | --- | --- |
| *Crassostrea gigas* | 63,340 | 17,784 | 2,008 | 12.6 | 3,301 | 286 |
| *Mytilus galloprovincialis* | 77,414 | 10,977 | 1,369 | 7.8 | 13,727 | 260 |
| *Mytilus coruscus* | 37,478 | 14,735 | 2,900 | 5.9 | 1,290 | 2,727 |
| *Dreissena polymorpha* | 68,018 | 13,316 | 2,603 | 4.5 | 1,632 | 1,194 |

**Table 6.** Statistical gene structure prediction for the blue mussel *M. chilensis* genome

|  | Type | Number | Average Transcript Length (bp) | Average CDS Length (bp) | Average Exons per Gene | Average Exon Length (bp) | Average Intron Length (bp) |
| --- | --- | --- | --- | --- | --- | --- | --- |
| De novo | Augustus | 83,110 | 10,628 | 1,423 | 5.74 | 1,423 | 1,941 |
|  | SNAP | 70,217 | 36,619 | 536 | 9.46 | 536 | 4,265 |
|  | GlimmerHMM | 39,700 | 1,973 | 753 | 2.85 | 753 | 658 |
|  | Geneid | 49,963 | 3,111 | 1,036 | 2.79 | 1,036 | 1,160 |
|  | Genscan | 83,110 | 10,628 | 1,423 | 5.74 | 1,423 | 1,941 |
|  | GeneMark | 81,256 | 11,018 | 1,301 | 5.82 | 1,301 | 270 |
| Homolog | GMAP | 250,833 | 16,107 | 370 | 3.10 | 1,649 | 6,891 |
|  | spaln | 136,264 | - | - | 7.80 | 179 | - |
| RNAseq | PASA | 18,056 | - | - | 2.05 | 504 | - |
| EVM |  | 63,943 | 6,137 | 1,133 | 4.54 | 1,133 | 1,414 |
| **Final set** |  | **34,530** | **6,531** | **1,377** | **4.92** | **1,377** | **1,316** |

**Table 7.** Statistics of Non-coding RNA annotation for the *M. chilensis* genome

| Type | Copy number | Average length(bp) | Total length(bp) | % of genome |
| --- | --- | --- | --- | --- |
| miRNA | 1365 | 92.8 | 126678 | 0.01% |
| rRNA | 143 | 118.58 | 16957 | 0.00% |
| sRNA | 275 | 83.68 | 23011 | 0.00% |
| snRNA | 99 | 106.71 | 10564 | 0.00% |
| snoRNA | 2267 | 89.38 | 202622 | 0.01% |
| tRNA | 516 | 77.44 | 39957 | 0.00% |
| tRNA pseudogenes | 123 | 72.96 | 8974 | 0.00% |
| All types | 4795 | 89.42 | 428763 | 0.02% |

**Table 8.** Statistics of gene function annotation for the *M. chilensis* genome

|  | Number | Percentage (%) |
| --- | --- | --- |
| Swissprot | 24,325 | 70.45 |
| Nr | 25,212 | 73.01 |
| Nt | 3,102 | 8.98 |
| KEGG | 22,425 | 64.94 |
| eggNOG | 23,181 | 80.57 |
| GO | 11,606 | 33.61 |
| Pfam | 60,887 | 176.33 |
| Annotated | 27,821 | 80.57 |
| Unannotated | 6,709 | 19.43 |
| Total | 34,530 | 100 |

### 3.4. Comparative genomics

Smooth-shelled blue mussels of the genus *Mytilus* represents a model group because of their cosmopolitan distribution, socioecological importance, and their intriguing evolutionary history. This taxon provides new insights into the process of speciation, and how the hybridization and introgression can be one of the biggest threats to global mussel’s biodiversity (Gardner, Oyarzun, Toro, Wenne, & Zbawicka, 2021). Survey of single nucleotide polymorphisms (SNPs) on Southern hemisphere blue mussels has provided a new layer for the understanding of their biology, taxonomy and phylogeography (Araneda, Larrain, Hecht, & Narum, 2016; Nunez-Acuna & Gallardo-Escarate, 2013). However, SNP markers cannot be applied as a single tool to evidence chromosome rearrangements events during the *Mytilus* evolution. Here, whole-genome sequencing in smooth-shelled blue mussels and relatives Bivalve species is a priority for global mussel aquaculture, biosecurity and conservation.

With the aim to explore genomic rearrangements in *Mytilus*, the reported reference genome for *M. coruscus* and *M. chilensis* were analyzed. Of the 34,530 predicted genes from the *M. chilensis* genome, 18,758 (54.32%) were found in syntenic collinear blocks after being compared with the *M. coruscus* genome (Fig. 3A). These syntenic blocks consisted of 671 alignments with a minimum of 5 genes per block. The number of alignments per chromosome ranged from 27 on chromosome 13 to 69 on chromosome 3. Chromosomes with the higher number of genes in collinear blocks were chromosomes 1, 4, and 6, with 1,227, 1,091, and 1,088 genes, respectively. Blocks with less than five genes or E-value < 1E-5 were discarded from this analysis. Most collinear blocks were located at the same pair of chromosomes between the two genomes. For example, *M. chilensis* Chr1 had only syntenic blocks with LG01 from *M. coruscus* in the same order. However, chromosomes 6 and 10 from *M. chilensis* had collinearity with chromosomes LG09 and LG02, respectively, in *M. coruscus* but were orientated as two inversed blocks per pair of chromosomes (red lines in Fig. 3A and Fig. 2S). The genes in these alignments from inversed blocks were extracted, blasted, and Gene Ontology terms were identified. Enrichment analyses from GO terms were obtained from Chr10 and LG09 inversed blocks and Chr6 and LG02 pair of chromosomes (Fig. 3B-C). Most of the molecular functions enriched GO terms in the Chr10 and LG09 pair were associated with heat shock protein (HSP) binding. In contrast, in the Chr6 and LG02 pair, most of the enriched GO terms were associated with the mitochondria and biological processes related to autophagy or regulation of the gene expression by epigenetic changes. Notably, chromosome rearrangements have been associated with adaptative genetic traits in marine organisms (Kess et al., 2020), where specific architectural proteins such as HSPs may have distinct roles in establishing 3D genome organization (L. Li et al., 2015).

**Figure 3.**
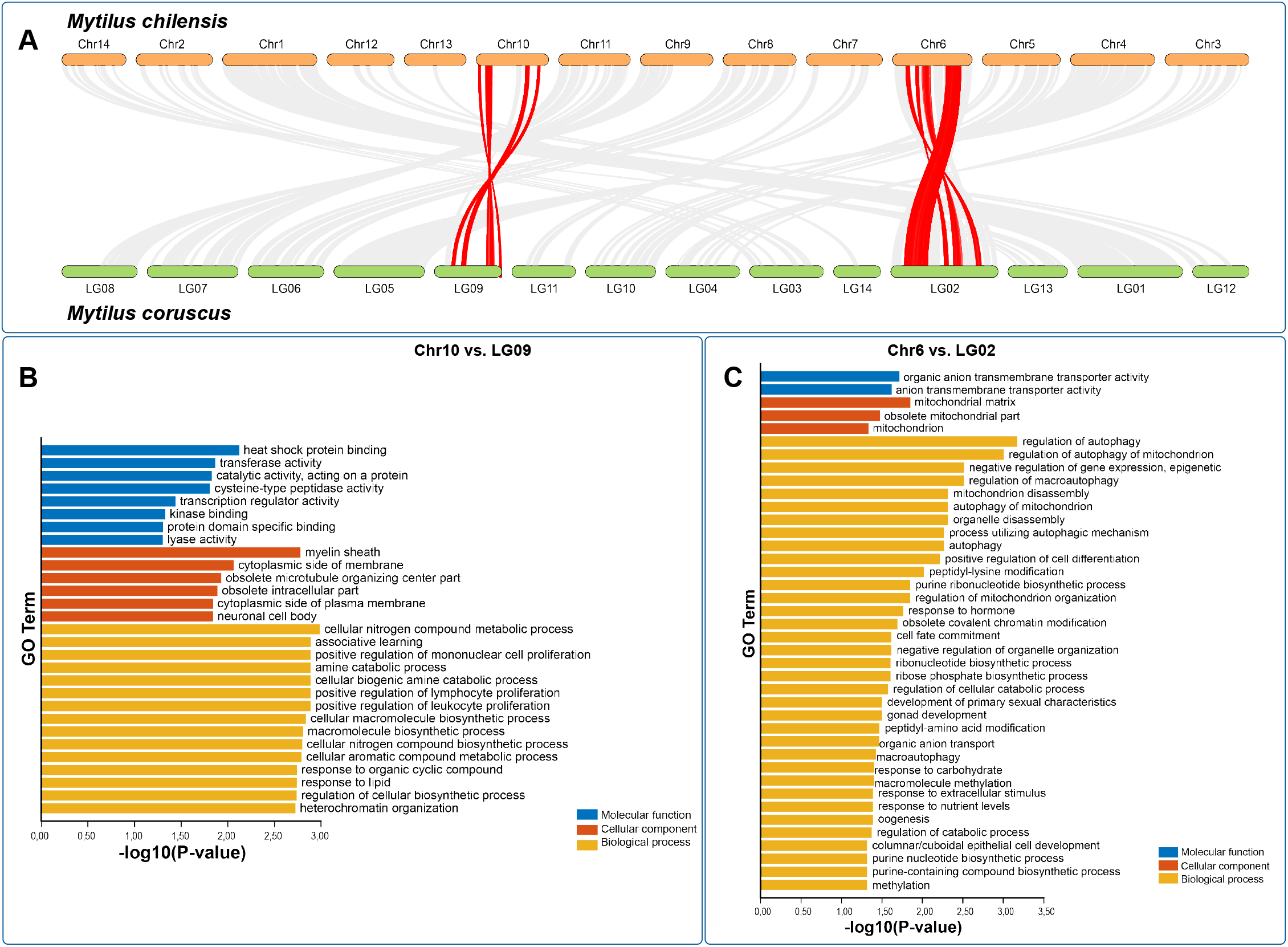
(A) Whole-genome macrosyntenic relationships between *M. chilensis* and *M. coruscus.* Orthologous relationships among mussel chromosomes are highlighted in grey and orthologous relationship between chr6 and LG02, and chr10 and LG09 are highlighted in red. (B-C) Gene Ontology term annotated for the syntenic chromosome regions identified by MCScanX for chr6 vs. LG02, and chr10 vs. LG09. The x-axis indicates the negative log_10_(q-value)

### 3.5. Comparative analysis of Steamer-like elements in Bivalvia

To explore the gene expansion of retrotransposon elements among representatives’ species from Bivalvia, we primarily characterized the Steamer-like elements (SLEs) in *M. chilensis* using the approach described by Arriagada et al. (2014). The analysis evidenced that the genome of *M. chilensis* contains five copies of SLEs distributed in chromosomes 1, 6, 7, 10, and 11. The alignment showed that all SLEs copies are flanked by two LTRs (5’ and 3’) containing the Gag-Pol ORFs and the domains annotated to Protease, Reverse Transcriptase, RNAaseH, and Integrase. Notably, an insertion composed of 12 nucleotides at position 933^934 was exclusively found in chromosomes 7 and 11. The translation for the inserted nucleotides suggests four amino acids, K, T, S, and H, in positive orientation. However, the translation evaluated in the reading frame (−1) evidenced a methionine localized before the RNAaseH coding gene (Fig. 4A).

**Figure 4.**
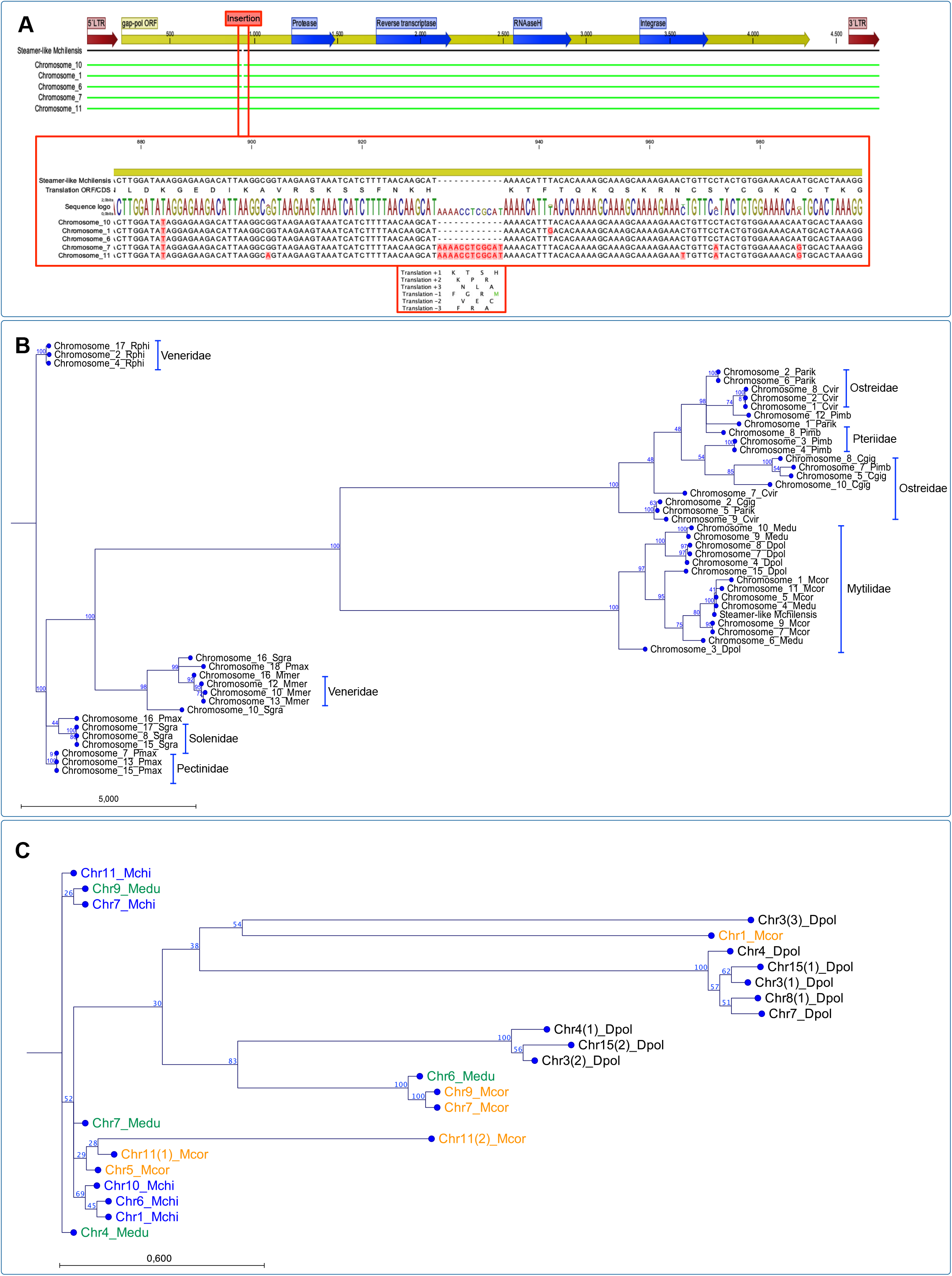
Molecular characterization of Steamer-like elements (SLE) in the *M. chilensis* genome and phylogenetic analysis using publicly available reference genomes assembled at chromosome-level for bivalve species. (A) Alignment of five SLE copies localized in the chromosomes 1, 6, 7, 10 and 11. All the SLE are flanked by two LTRs (5’ and 3’) containing the Gag-Pol ORFs and the domains annotated for Protease, Reverse Transcriptase, RNAaseH and Integrase. An insertion composed by 12 nucleotides (933^934 position) was found in the chromosomes 7 and 11. The detailed alignment and the translation for the nucleotides inserted are highlighted in the red box. (B) Maximum likelihood (ML) phylogenetic tree of nucleotide sequences from SLEs found in eleven reference genomes for Bivalvia. Colored chromosomes and numbers indicate the SLE genome localization and the bivalve species, respectively. The species analyzed were: Veneridae (pink), *Ruditapes philippinarum* (Rphi) and *Mercenaria mercenaria* (Mmer). Solenidae (grey) *Solen grandis* (Sgra). Pectinidae (blue), *Pecten maximus* (Pmax). Ostreidae (green) *Crassostrea gigas* (Cgig), *C. virginica* (Cvir) and *C. ariakensis* (Caria). Pteriidae (blue light), *Pinctada imbricata* (Pimb). Mytilidae (red), *Mytilus coruscus* (Mcor), *Mytilus edulis* (Medu) and *Dreissena polymorpha* (Dpol). (C) ML analysis of SLEs identified in *M. edulis* (orange), *M. coruscus* (red), *D. polymorpha* (purple) and *M. chilensis* (blue) chromosomes.

Furthermore, the phylogenetic analysis using publicly available reference genomes assembled at chromosome level for eleven bivalve species using Maximum Likelihood (ML) revealed six chromosomes cluster composed of bivalves belonging to the families Veneridae, Solenidae, Pectinidae, Ostreidae, Pteriidae, and Mytilidae (Fig. 4B). The phylogenetic reconstruction rooted SLEs found in three chromosomes (2, 4 and 17) from *R. philippinarum.* The other Veneridae member, *M. mercenaria* showed a cluster of four chromosomes (10, 12, 13, and 16) and related to two chromosomes of *S. Grandis* (10 and 16). This last species formed a unique cluster composed of three chromosomes (8, 15, and 17), similar to *P. maximus*, with three chromosomes. Concerning the mussel and oyster genomes assembled at the chromosome level, the phylogenic analysis revealed two main clusters composed of species belonging to Ostreida and Mytilidae, where the first taxon was comprised of the Ostreidae and Pteriidae families. Herein, one cluster was rooted with three SLE sequences from *C. virginica*, *C. gigas*, and *C. ariakensis* located on chromosomes 9, 2, and 5, respectively. The second major cluster was composed of SLEs annotated in chromosomes from Ostreidae and Pteriidae, where *C. virginica* chromosomes were closely related to *P. imbricata*. The third cluster was observed containing three SLE sequences from *C. virginica* and *C. ariakensis*; chromosomes 1, 2, 8, and 1, 2, and 6, respectively (Fig. 4B). The analysis of the Mytilidae family revealed two primary clusters comprised of SLE located in chromosomes from *M. edulis and D. polymorpha*, and *M. coruscus*, *M. chilensis*, respectively (Fig. 4B). This last cluster grouped five chromosomes from *M. coruscus* (Chr. 1, 4, 5, 7 and 11), and two from *M. edulis* (Chr. 4 and 6). The Steamer-like sequenced characterized for *M. chilensis* was also observed in this cluster. Finally, a detailed analysis of the three mussel species reported with genome assemblies at the chromosome level was conducted (Fig. 4C). Notably, a rooted cluster comprising chromosomes 7, 11, 9 for *M. couscous*, and 4 and 6 for *M. edulis* were closely related. Herein, two primary clusters of SLEs located in chromosomes from *M. chilensis*, *M. eduli*s, and *M. coruscus* were observed. The analysis suggested that the SLEs identified on *M. chilensis* chromosomes are closely related to the SLEs annotated on chromosomes 9 and 4 in *M. edulis*; meanwhile, the SLEs located in chromosomes 1, 5, and 11 in *M. coruscus* were also identified in the same chromosome cluster. The second main cluster observed was comprised exclusively of SLEs annotated in *D. polymorpha* chromosomes, except by the SLEs copies identified in chromosomes 9 and 10 of *M. edulis*. Interestingly, the SLEs annotated in chromosome 9 from *M. edulis* are shared among the three primary clusters analyzed, suggesting putative translocation gene events in Mytilidae. Overall, the phylogenic relationships of SLEs revealed that the reported bivalve genomes comprise between 3 to 6 loci. A lower number of SLEs was found in Solenidae, Pectinidae, and Veneridae, followed by Mytilidae. A higher number of SLE loci was observed in genomes belonging to the Ostreida order. As far as we know, the evolution of the bivalve chromosomes has mainly been studied using cytogenetical techniques combining molecular probes on candidate genes to detect genome rearrangements that drive the speciation process (Garcia-Souto, Perez-Garcia, Kendall, & Pasantes, 2016; Gonzalez-Tizon, Martinez-Lage, Rego, Ausio, & Mendez, 2000; Y. Wang & Guo, 2004). However, the availability of reference genomes assembled at the chromosome level opens new perspectives to explore the molecular evolution at several taxonomic orders through gene collinearity analysis. The study by Yang et al. (2021) highlighted putative chromosome rearrangements among the king scallop *Pecten maximus*, the blood clam *Scapharca broughtonii*, the hard-shelled mussel *Mytilus coruscus*, the pearl oyster *Pinctada martensii*, and the Pacific oyster *Crassostrea gigas* genomes. Notably, the chromosome synteny illustrated that large-scale rearrangements are common events between scallop and oysters but scarce between scallop and mussel genomes. The reported evidence suggested that almost all the chromosome rearrangements between the mussel and the oyster genomes are different, implicating independent chromosome fusion events. The SLEs loci identified in all the genomes analyzed in the current study suggest that SLEs are relatively conserved in chromosome position for some taxa. For instance, the SLEs loci in Veneridae, Pectinidae, and Solenidae appear to be associated with chromosomes 10, 13, 12, and 16. This sharing characteristic can reflect common genetic events during the evolution of these taxonomical groups. Similarly, the Ostreidae and Mytilidae families share SLEs loci annotated to chromosomes 1, 2, 8, and 10. The detailed analysis of SLEs in Mytilidae evidence that transposon identified in *M. chilensis* was shared between *M. edulis* and *M. coruscus*, where SLEs in *D. polymorpha* appear to be more phylogenetically distant than *Mytilus* species. Interestingly, the mutation identified on the SLEs localized in the *M. chilensis* genome (insertion of twelve nucleotides), specifically on chromosomes 7 and 11, was shared with the SLE annotated on chromosome 9 in *M. edulis*. This cumulative evidence reveals diverse chromosome rearrangements, reflecting a complex evolutionary history of bivalve chromosomes.

### 3.6. The marine environment of *M. chilensis* populations

Temporal and spatial variability of sea surface temperature (SST) around Chiloé island and at Yaldad and Cochamó were analyzed over the past two decades. The oceanographic variability for the location studied was analyzed from remote sensing data and in situ measurements (Fig. 5A-D). Here, the daily time series of SST extracted from satellite-derived data for both sites evidenced high surface temperature variability between Yaldad and Cochamó, where this last location was constantly higher thought the year (Fig 3S). Notably, the monthly medians computed from the SST time series showed that the main differences were observed during the austral summer (from December to March. During the winter season, the oceanographic variability was less pronounced, showing the temperatures between 13°c and 10°C from April to July (Fig. 5C). Furthermore, *in situ* data were collected from June 2017 to May 2018 (exact time where the specimens were collected), exhibiting significant differences in both location for temperature and salinity among 0 and 20 meters of depth. Interestingly, the oceanographic survey revealed a pronounced vertical stratification with higher temperatures and lower salinity in Cochamó compared with Yaldad (Fig. 5D). These observations support the idea that there are two oceanographically different zones in the inner sea of Chiloé Island. In this northern area, mussels from Cochamó and Yaldad were sampled for the current study. Taken together, we can hypothesize that the mussels inhabiting Cochamó are significantly more exposed to environmental stress than the Yaldad mussel population. To date, there are few studies exploring how population genetic variation is related to, or caused by, the marine environmental variation in mussel populations. Notably, a study conducted by Wenne et al. (2022) examined the genetic differentiation of native populations of *M. galloprovincialis* throughout its entire geographic range in the Mediterranean Sea, the Black Sea and the Sea of Azov using 53 SNP loci. The results indicated that seven of the 13 environmental variables explained significant variation in population-specific SNP locus allele frequencies. These seven variables explained a total of 75% of the variation in the SNP dataset, suggesting that there is a complex mix of environmental variables that contribute to genetic variation of *M. galloprovincialis* populations in the Mediterranean Sea.

**Figure 5.**
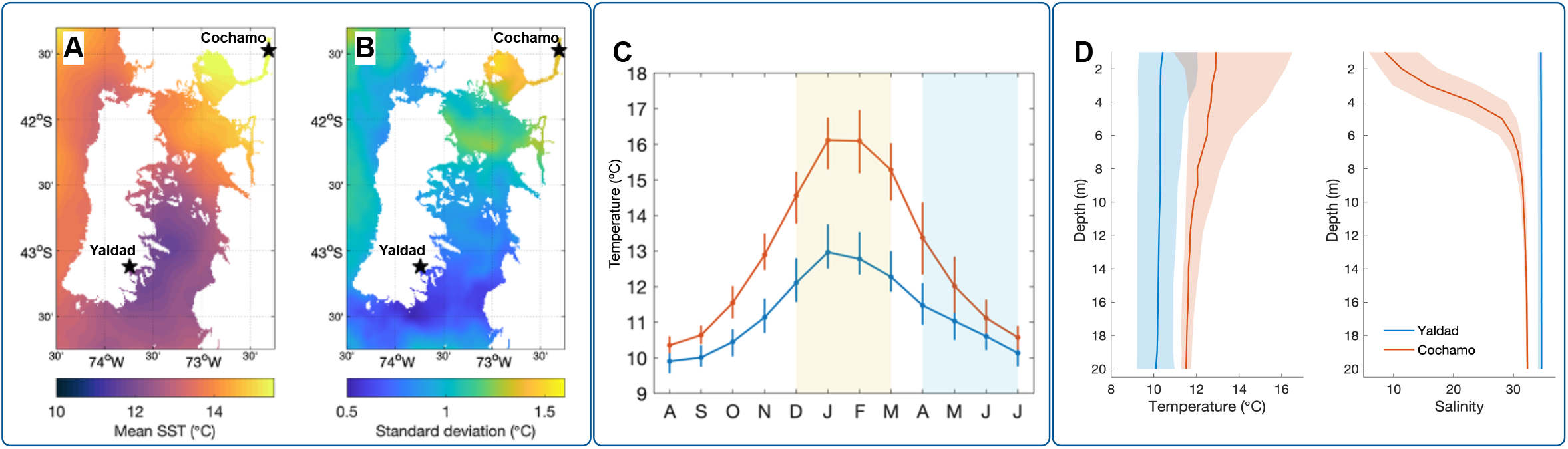
Temporal and spatial variability around Chiloé island, and at sites Yaldad and Cochamo, over the past two decades. (A-B) Maps correspond to mean SST computed for December 2017 – March 2018 and the corresponding standard deviations. Stars indicate the location of sampling sites. (C) Monthly medians computed from data in (A), showing the first and third quartiles as error bars; note that the sequence of months shown on the x-axis begins in August and ends in July. Shaded areas indicate summer (yellow) and fall-winter (blue) periods. (D) *In situ* measurements for temperature (°C) and salinity (PSU) from 0 to 20 meters of depth. The blue and red lines for (C) and (D) represent the data collected from Yaldad and Cochamo, respectively.

### 3.7. Whole-genome transcript expression analysis in two *M. chilensis* populations

The transcriptome profiling among mussels collected from Yaldad and Cochamó evidenced three primary transcriptional clusters, where gene cluster 1 was highly expressed in the gills of mussels exposed to the Yaldad marine conditions; meanwhile, gene clusters 2 and 3 were highly expressed from individuals collected in Cochamó or mussel exposed to estuarine conditions (Fig. 6A). The RNA-seq analysis was performed with the mRNA sequences annotated on the *M. chilensis* genome. Herein, it is essential to note that in mussel species, specifically in *M. galloprovincialis*, the phenomenon of Presence-Absence Variation (PAV) has been described. This fact means that PAVs can bias the analyses of transcriptome profiles in the studied mussel populations. We previously conducted a *de novo* assembling for the RNA-data sets sequenced from Yaldad and Cochamó populations (data not shown). The results evidenced that the completeness of the gene set annotated in the *M. chilensis* genome did not show statistical differences between both mussel populations. The evaluation of differentially expressed genes (DEGs) showed that the main factor of differences in the amount of DEGs was due to the population more than the replicates assessed (Fig. 6B). The proportion of DEGs evaluated among the gene clusters revealed that the cluster 1, highly expressed in Yaldad, accounted the 78.85% of the total DEGs analyzed. Clusters 2 and 3 are primarily characterized by high transcription values in the Cochamó population, evidenced by 7.32% and 13.82% of DEGs, respectively. The total number of DEGs analyzed was 1,570 (Fig. 6C). Notably, the fold-changes values estimated among the replicates and populations revealed high values in the gene transcriptional cluster 1, compared with clusters 2 and 3 where the fold-changes values were significantly lower (Fig. 6D). The functional analysis showed that the cluster 1 was enriched by GO terms related with protein modification processes, programmed cell death, immune system processes, defense response, cell differentiation and anatomical structure development (Fig. 6E). The clusters 2 and 3 were less enriched, revealing significant GO terms for transmembrane transport, reproductive processes, protein-containing complex assembly, microtube-based movement, cytoskeleton organization, chromatin organization and metabolic process (Fig. 6E).

**Figure 6.**
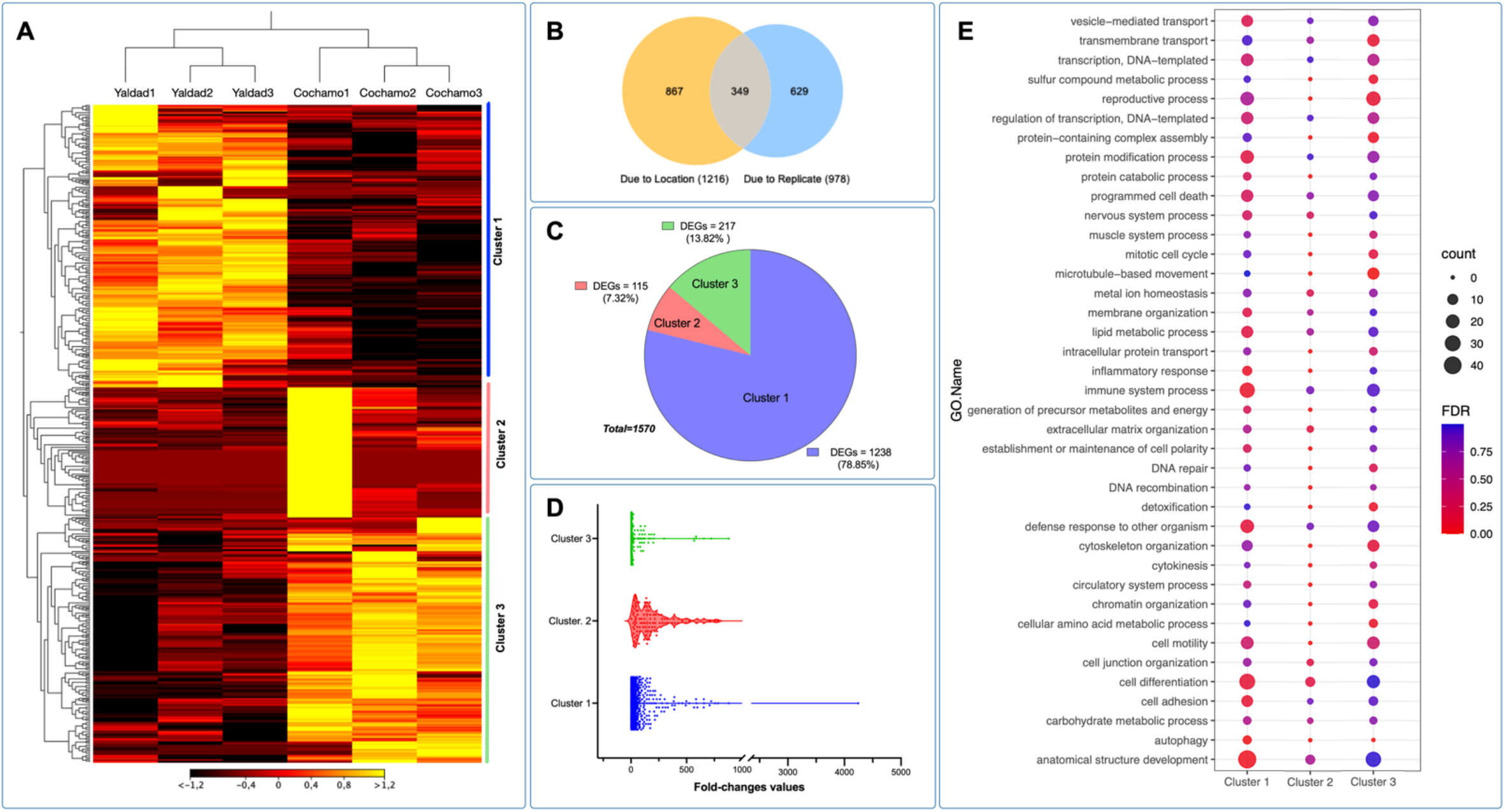
Population-specific transcriptome analysis in the blue mussel *M. chilensis*. (A) Transcriptome patterns of coding genes analyzed in gills from Yaldad and Cochamo population. Three replicates were evaluated from each experimental group. The heatmap was based on Transcripts Per Million (TPM) calculation and hierarchical clustering on Manhattan distances with average linkage. Yellow colors mean upregulated coding genes, black represents downregulated genes. (B) Venn diagram showing shared and unique genes expressed among the locations and replicates. (C) Pie chart showing number of differentially expressed genes (DEGs) annotated for expression cluster 1, 2 and 3 between Yaldad and Cochamo populations. (D) Fold-changes values observed for DEGs identified in each cluster evaluated. (E) GO enrichment of cluster-specific genes (P-value<10–16;|fold-change|>5) annotated for key biological processes differentially expressed. The y-axis indicates the GO term, and the x-axis indicates the negative log_10_(q-value). The color bar indicates the enriched factor. The bubble size indicates the number of GO terms.

The cluster gene expression analysis was used to identify genetic polymorphisms annotated in differentially expressed genes (DEGs) between Yaldad and Cochamó mussel populations. The evaluation of DEGs was performed by cluster transcriptome analysis displayed using a Circos plot to visualize specific loci where DEGs were highly transcribed. The fold-change values calculated showed high levels of transcription in clusters 1 and 2 through all chromosomes scanned (see red dots in Fig. 7A). Congruently with the previous RNA-seq results in this study, the highest fold-change values were observed in DGEs annotated in cluster 1 (Yaldad population). On the contrary, cluster 3 showed a small number of DEGs with high fold-changes values. Notably, the DEGs localized on the *M. chilensis* chromosomes evidenced specific-transcriptome patterns where some genes are differentially and spatially expressed through the genome of mussels exposed to the Yaldad and Cochamó environment conditions. The synteny analysis for DEGs and differentially localized in the chromosomes showed a marked pattern of syntenic relationships among chromosomes 5, 7, and 12 for cluster 1; meanwhile, the synteny observed for the DEGs annotated in clusters 2, and 3 revealed a wide distribution along the *M. chilensis* genome (Fig. 7A). Interestingly, the analysis carried out to detect macro-genome mutation in gene families between Yaldad and Cochamó population evidenced a similar number of dispersed genes, suggesting that those might arise from transposition. Tandem genes or repeatedly duplicated were observed with a low proportion in cluster 3 (Cochamó); meanwhile, the proximal genes showed a similar proportion to cluster 1 (Yaldad). These results might suggest small-scale transposition or duplication/insertion events. An interesting finding was observed for whole genome duplication (WGD). The primary proportion was evidenced in cluster 3 (Cochamó), compared with clusters 1 and 2 from the Yaldad population (Fig. 7B). Furthermore, the bioinformatic analysis conducted for detecting amino acid changes (AAC) in DEGs showed that 38% of non-synonymous AAC were identified in mussels collected from Yaldad. Contrary, the main proportion of synonymous AAC was detected in mussels exposed to the Cochamó’s estuarine conditions (Fig. 7C). Notably, the analysis performed for DEGs annotated in cluster 2 did not show non-synonymous and synonymous AAC in mussels collected from Yaldad. Finally, the evaluation of the zygosity proportion estimated for each mussel population evidenced an inverse pattern between both populations. The Yaldad cluster was higher in the homozygous proportion compared with Cochamó, where AAC heterozygous were detected in a higher proportion (Fig. 7D).

**Figure 7.**
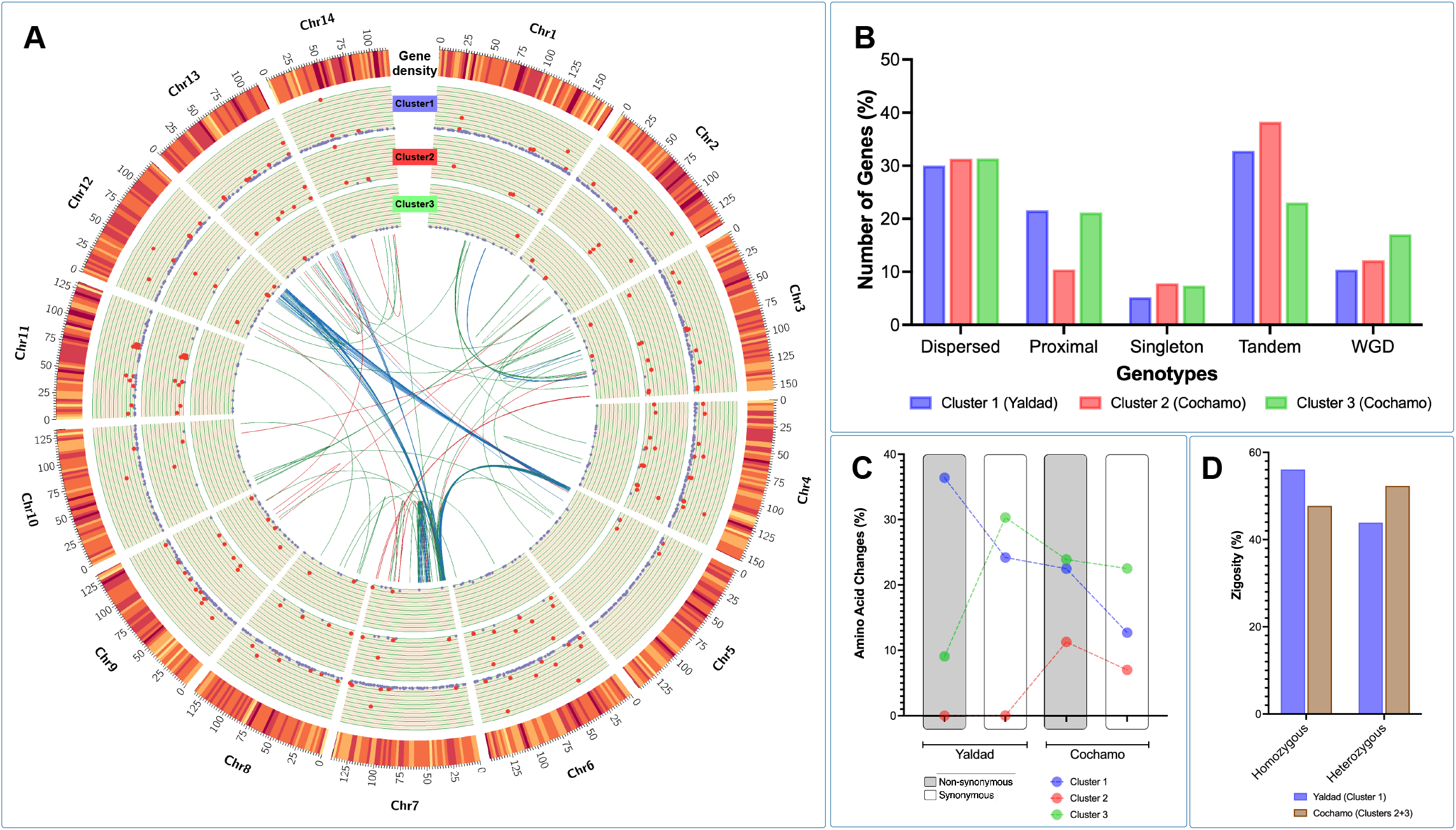
Genetic polymorphisms annotated in differentially expressed genes (DEGs) analyzed between Yaldad and Cochamo mussel populations. The evaluation of DEGs was performed by cluster transcriptome analysis. (A) Circos plot showing DEGs identified in the three analyzed clusters. From outer to the inner circle: Gene density, DEGs cluster 1, DEGs cluster 2, DEG cluster 3 and syntenic relationships between DGEs (each color line represents the cluster analyzed). Red dots represents DEGs with fold-changes values >|100| and purple dots represents fold-changes values <|10|. (B) Genotypes of DEGs identified in *M. chilensis* populations according to the transcription cluster analysis. Singleton means that the gene is single-copy, which should not be the type of members of gene families. Dispersed means that the gene might arise from transposition. Tandem means that genes were repeatedly duplicated. Proximal means that the gene might arise from small-scale transposition or arise from tandem duplication and insertion of some other genes and whole genome duplication (WGD) means that the gene might arise from chromosome duplication region. The analysis was carried out using *MCScanX*. (C) Amino acid changes proportions (%) between Yaldad and Cochamo populations. For each mussel population, the non-synonymous and synonymous were annotated for the DEGs selected according to the cluster analysis. (D) Zygosity proportion (%) estimated for each mussel population. Cluster 1 (Yaldad) is represented by the blue bars and the cluster 2 and 3 (Cochamo) is displayed in brown color bars.

To explore the transcriptome signatures between Yaldad and Cochamó mussel populations, we applied the genome chromosome expression (CGE) approach to test differences among tissues and individuals through the *M. chilensis* genome. The CGE analysis revealed high differences among chromosome regions, where the gills tissue was more modulated than the mantle tissue (Fig. 8A). Interestingly, there are some levels of congruence among the CGE annotated for both mussel populations. Using this finding, we conducted a gene ontology enrichment analysis from genes identified by CGE analysis. The results evidenced that gills transcriptomes displayed functional processes associated with transmembrane transport, protein catabolic, nervous system, and metal homeostasis. Notably, immune system process GO terms were highly enriched in gills. Moreover, the chromosome region differentially expressed in mantle tissue revealed that the reproductive process, protein modification process, cell differentiation, anatomical structure development, and gene silencing by RNA were mainly annotated (Fig. 8B). Taken together, the results reported in this study are highly congruent with the previous study conducted by Yévenes et al., (2022) through the transcriptome responses of *M. chilensis* collected in ecologically different farm-impacted seedbeds.

**Figure 8.**
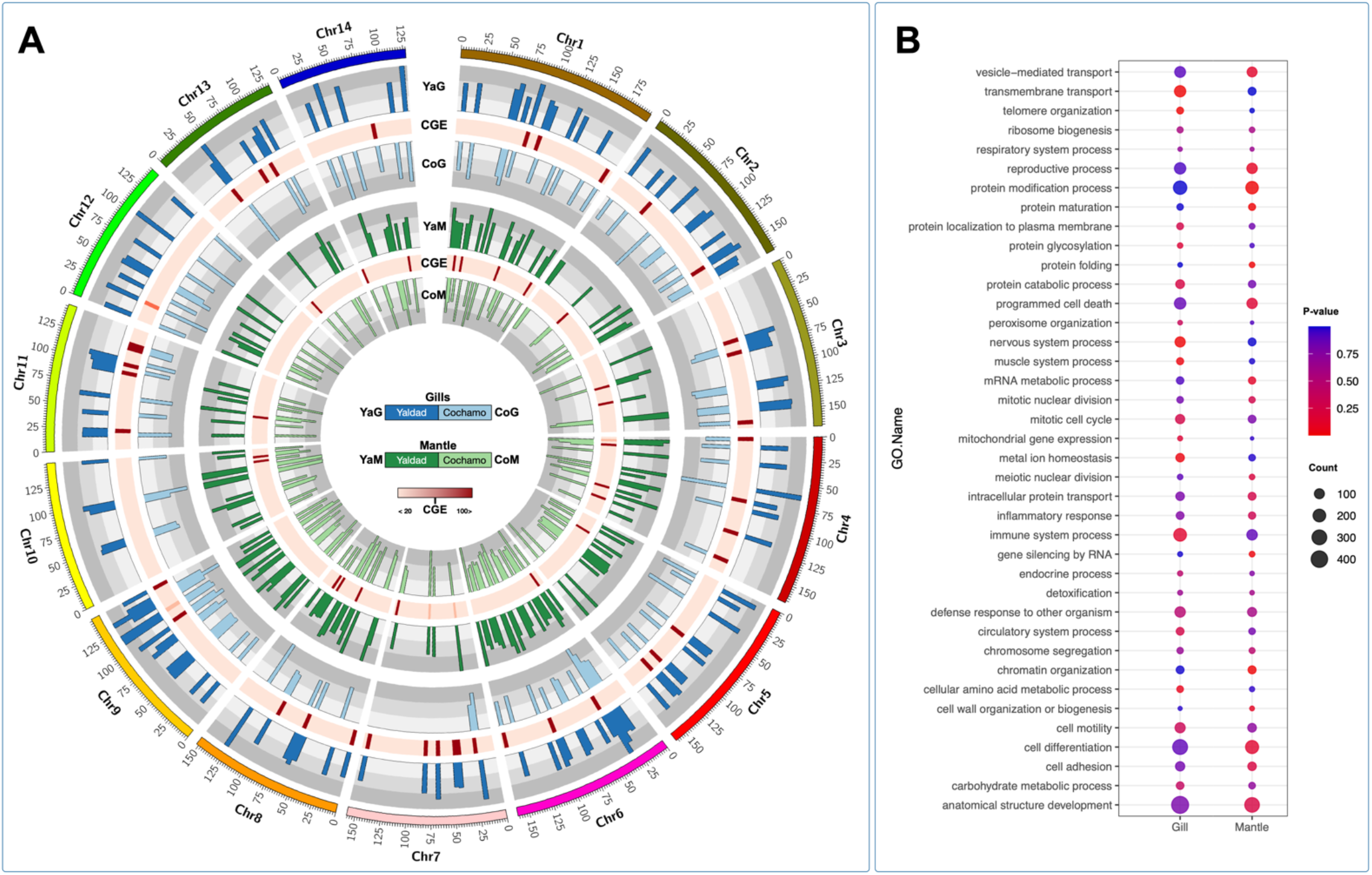
Chromosome-genome transcription in *M. chilensis* tissues collected from Yaldad and Cochamo populations. A) Threshold of gene expression from gills and mantle were mapped and compared on chromosomes regions from the two analyzed mussel populations. Transcriptional differences among locations (Yaldad/Cochamo) and tissues (Gills/Mantle) were used to estimate the CGE index. Heatmap in red shows the variation of gene expression from high to low differences. (B) GO enrichment of tissue-specific genes (P-value<10–16;|fold-change|>5) annotated for key biological processes differentially expressed. The y-axis indicates the GO term, and the x-axis indicates the negative log_10_(q-value). The color bar indicates the enriched factor. The bubble size indicates the number of GO terms.

The cumulative findings archived in this study strongly suggest that the immune system was primarily modulated between mussels exposed to Yaldad and Cochamó environmental conditions. With the aim to explore the transcription profiling of immune-related genes, we selected two KEGG pathways annotated in the *M. chilensis* genome (Fig. 9). Herein, TOLL-like receptor signaling pathway and apoptosis were analyzed in terms of transcription activity and single nucleotide variation (SNV) between mussel populations. Notably, a non-synonymous SNV was detected on the TLR2 gene (28T>G) in individuals collected from the Yaldad population. The translation evidenced an amino acid change from Phenylalanine to Valine at position 10 in the ORF (Phe10Val) (Fig. 9A). The analysis also evidenced SNV on the genes such as AKT and TAB1, where no amino acid changes were detected. The transcriptome profiling for the TLR pathway evidenced a high modulation of genes such as TLR3, AKT, TRAFF6, FADD, IRAK4, and RAC1 in mussels collected from Yaldad. Interestingly, mitogen-activated protein kinase kinases (MAP2K and MAPK1) and c-Jun N-terminal kinase (JNK) were differentially expressed, suggesting putative roles related to stress signaling pathways (Fig. 9B). Furthermore, the Apoptosis pathway revealed two SNV localized in *eukaryotic initiation factor 2 alpha* (eiF2⍺) and *inhibitor of apoptosis* (IAP) in mussel sampled from Yaldad and Cochamó population, respectively (Fig. 9C). The 2613delG in *eiF2⍺* gene produces a frameshift at the Thr872, meanwhile the 968_970delCTC localized in the IAP gene produces a deletion of proline at the position 323. The transcriptome profiling of apoptosis-related genes showed a conspicuous differentiation between gills and mantle tissue, where three primary gene expression clusters were identified (Fig. 9D). Notably, in gills tissue, genes such as P53, ERK, TP53, PARP2, and JNK were highly expressed. The gene expression analysis in mantle tissue evidenced high transcriptional activity in genes related to the intrinsic (mitochondria-mediated) pathway, such as *the B-cell lymphoma* (BCL) gene and the second mitochondria-derived activator of caspase (DIABLO) gene. Further functional studies will be conducted to validate the association between single nucleotide polymorphisms and the *fitness* traits observed or how the translocation process associated with the aquaculture activity can evolve in the loss of locally adapted alleles. Interestingly, a recent transplant experiment reported by Jahnsen-Guzmán et al. (2021) demonstrated that *M. chilensis* individuals are adapted to the subtidal environment (4 m depth), as they exhibit significantly higher fitness (growth and calcification rates) than those transferred to the intertidal environment (1 m depth), which showed increased metabolic stress. Herein, mussel lives in an extreme environmental variability, where their ability to cope with perturbations is based on the genome architecture to build plastic and adaptive responses.

**Figure 9.**
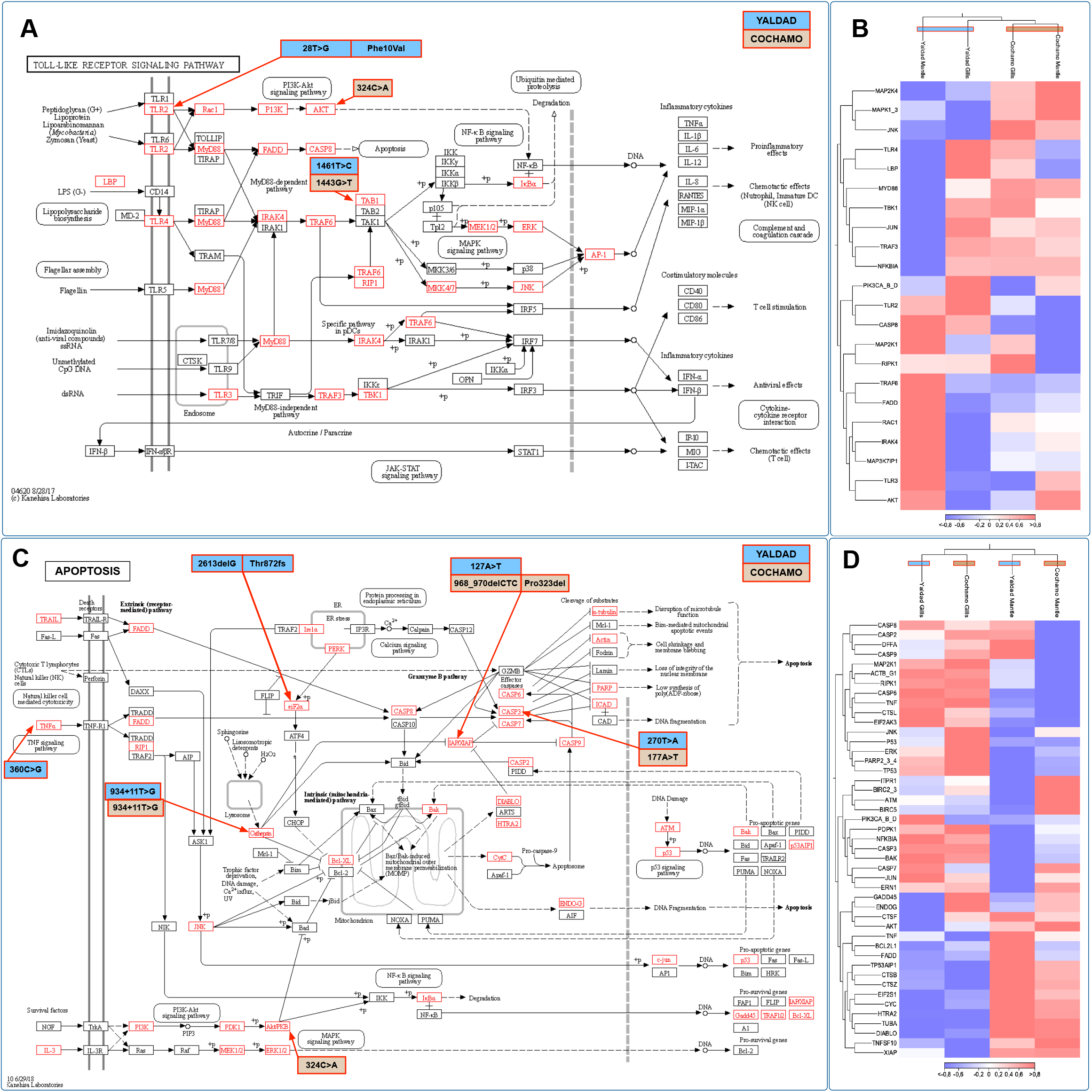
Transcriptome response of the immune system by KEGG pathways analysis and single mutation variant detection between *M. chilensis* populations. (A) TOLL-like receptor (TLR) and (C) Apoptosis signaling pathways comparisons between mussels from Yaldad and Cochamo. Single nucleotide polymorphisms and amino acid changes are shown in blue and brown boxes according to the mussel population. Identified gene families on the KEGG pathways are marked with red in the *M. chilensis* genome. (B) and (D) Transcriptome patterns of coding genes analyzed in gills and mantle from Yaldad and Cochamo population for TLR and Apoptosis-related genes. Three replicates were evaluated from each experimental group. The heatmap was based on Transcripts Per Million (TPM) calculation and hierarchical clustering on Manhattan distances with average linkage. Red colors mean upregulated coding genes, and blue colors represent downregulated genes.

In summary, we have generated the first chromosome-level assembly of the native blue mussel *Mytilus chilensis* genome. This genomic resource was used to identify genome signatures putatively related to the local adaptation process in the mussel population inhabiting contrasting marine environments. Collectively, the identification of putative mutations associated with immune and metabolic-related genes suggests that mussel populations facing highly variable environments display a genomic adaptation to reduce the number of genes and their transcriptional activity. This evolutionary strategy can suggest that the expression of those genes has evolved a degree of “frontloading” that potentially pre-adapts the mussel populations to frequent heat and salinity stress, contributing to their physiological tolerance and *fitness*. We believe that the generated genomic resource will be instrumental for future research on population genomics informing management and sustainable strategies for the Chilean mussel aquaculture.

## Author contributions

CGE designed and supervised the study. VV, GNA, DVM, JT, PO, and MY collected and prepared the mussel samples. CGE, CGE, VVM, DVM, GNA, and FT analyzed all sequencing and oceanographic data. CGE, GG, AF, BN, SR and MG wrote the manuscript with the other authors’ help. GA and CGE sequenced and analyzed the Steamer-like elements in *M. chilensis*. All authors revised the draft and approved the final manuscript.

## ACKNOWLEDGMENTS

This study was funded by FONDAP #15110027, FONDECYT #1210852 and #1130852, granted by ANID-Chile.

## DATA AVAILABILITY AND BENEFITS-SHARING STATEMENT

Data availability statement:

The Mytilus chilensis whole genome sequencing data supporting this study’s findings are available in NCBI under BioProject PRJNA861856. The sequencing data supporting this study’s findings are available in SRA at SRR20966976, SRR20593343 and SRP261955. Benefits from this research accrue from sharing our data and results on public databases as described above. The assembled genome and the genome annotation results were deposited in the Figshare database (https://doi.org/10.6084/m9.figshare.20995963.v1).

